# Coral bleaching susceptibility is predictive of subsequent mortality within but not between coral species

**DOI:** 10.1101/2019.12.17.880161

**Authors:** Shayle Matsuda, Ariana Huffmyer, Elizabeth A. Lenz, Jen Davidson, Joshua Hancock, Ariana Przybylowski, Teegan Innis, Ruth D. Gates, Katie L. Barott

## Abstract

Marine heat waves instigated by anthropogenic climate change are causing increasingly frequent and severe coral bleaching events that often lead to widespread coral mortality. While community-wide increases in coral mortality following bleaching events have been documented on reefs around the world, the ecological consequences for conspecific individual colonies exhibiting contrasting phenotypes during thermal stress (e.g. bleached vs. not bleached) are not well understood. Here we describe the ecological outcomes of colonies of the two dominant reef-building coral species in Kāne□ohe Bay, Hawai□i, *Montipora capitata* and *Porites compressa*, that exhibited either a bleaching susceptible phenotype (bleached) or resistant phenotype (non-bleached) following the second of two consecutive coral bleaching events in Hawai□i in 2015. Conspecific pairs of adjacent bleaching susceptible vs. resistant corals were tagged on patch reefs in two regions of Kāne□ohe Bay with different seawater residence times and terrestrial influence. The ecological consequences (symbiont recovery and mortality) were monitored for two years following the peak of the bleaching event. Bleaching susceptible corals suffered higher partial mortality than bleaching resistant corals of the same species in the first 6 months following thermal stress. Surprisingly, *P. compressa* had greater resilience following bleaching (faster pigment recovery and lower post-bleaching mortality) than *M. capitata*, despite having less resistance to bleaching (higher bleaching prevalence and severity). These differences indicate that bleaching susceptibility of a species is not always a good predictor of mortality following a bleaching event. By tracking the fate of individual colonies of resistant and susceptible phenotypes, contrasting ecological consequences of thermal stress were revealed that were undetectable at the population level. Furthermore, this approach revealed individuals that underwent particularly rapid recovery from mortality, including some colonies over a meter in diameter that recovered all live tissue cover from >60% partial mortality within just one year. These coral pairs continue to be maintained and monitored in the field, serving as a “living library” for future investigations on the ecology and physiology of coral bleaching.

## Introduction

Ocean warming due to anthropogenic climate change has caused an increase in the frequency and severity of coral bleaching, a visually striking stress response where the coral host loses its endosymbiotic algae (dinoflagellates of the family Symbiodiniaceae) thus revealing the white coral skeleton beneath the translucent animal tissue (Gates *et al.*, 1992; Putnam *et al.*, 2017). In this nutritional symbiosis, corals are obligate partners that require the algae for the majority of their energy (Muscatine and Porter, 1977; Muller-Parker *et al.*, 2015). Because of coral dependence on this partnership, prolonged heat waves that cause sustained coral bleaching can lead to depletion of the host’s energy supply and reserves (Grottoli *et al.*, 2004; Rodrigues and Grottoli, 2007; Imbs and Yakovleva, 2012; Wall *et al.*, 2019). This stress can elicit a variety of sublethal effects, including declines in growth and reproduction (Ward *et al.*, 2000; Baird and Marshall, 2002; Baker *et al.*, 2008; Hughes *et al.*, 2019), and at worst can result in widespread coral mortality (Loya *et al.*, 2001; McClanahan, 2004; Baker *et al.*, 2008; Eakin *et al.*, 2010; Hughes *et al.*, 2017; Sully *et al.*, 2019). The loss of live coral cover is often followed by rapid erosion of the structural framework of the reef (Hughes, Kerry, *et al.*, 2018; Fordyce *et al.*, 2019; Leggat *et al.*, 2019), reducing habitat complexity (Magel *et al.*, 2019) and negatively impacting the diversity of the broader reef community (e.g. fish (Pratchett *et al.*, 2011; Darling *et al.*, 2017; Richardson *et al.*, 2018). Coral mortality following bleaching can also alter the structure of the coral community itself, as bleaching-susceptible species are lost from the community (Loya *et al.*, 2001; McClanahan, 2004; Baker *et al.*, 2008; Bahr *et al.*, 2017; Hughes *et al.*, 2017) while stress-tolerant species remain (Edmunds, 2018; Hughes, Kerry, *et al.*, 2018) and “weedy” genera that are better suited for rapid recovery following bleaching become dominant (Darling *et al.*, 2012; Edmunds, 2018). These changes in community composition alter the ecological function of the reef (Alvarez-Filip *et al.*, 2013), which alongside the structural degradation following bleaching leads to declines in ecosystem goods and services ranging from fisheries production to coastal protection and tourism (Munday et al. 2008).

The susceptibility of a reef system to coral bleaching during a marine heat wave depends on a combination of physical and biological factors (Swain *et al.*, 2016). Differences in the local microenvironment (e.g. flow, turbidity, surface reflectance, internal waves, etc.) can ameliorate or exacerbate the magnitude of thermal stress and lead to differential bleaching and mortality on nearby reefs (Anthony *et al.*, 2007; Wyatt *et al.*, 2019). Local temperature dynamics can also influence coral bleaching by altering the physiological tolerance of coral populations to heat stress. For example, corals exposed to high diel temperature variability often have higher thermal tolerance and greater resistance to bleaching than nearby conspecifics in more stable regimes (Putnam and Edmunds, 2011; Palumbi *et al.*, 2014; Schoepf *et al.*, 2015, 2019; Barshis *et al.*, 2018). These data indicate that environmental history of individuals and populations plays a significant role in determining coral responses to stress, and is likely driven by a combination of both acclimatization (Bay *et al.*, 2013; Bay and Palumbi, 2015; Kenkel and Matz, 2016) and adaptation (Barshis *et al.*, 2013, 2018; Palumbi *et al.*, 2014; Matz *et al.*, 2018). On a global scale, there is recent evidence that the temperature threshold for coral bleaching has risen in correspondence with global warming, suggesting widespread acclimatization and/or adaptation (Coles *et al.*, 2018; DeCarlo *et al.*, 2019). Alternatively, differences in the composition of coral communities between reefs can also influence bleaching extent due to species-specific differences in thermal tolerance (Rowan *et al.*, 1997; Berkelmans and Van Oppen, 2006; Sampayo *et al.*, 2008; Swain *et al.*, 2016; Edmunds, 2018). Indeed, the global loss of many thermally sensitive coral species from reef communities (e.g. (Loya *et al.*, 2001; McClanahan, 2004; Hughes, Kerry, *et al.*, 2018; Kim *et al.*, 2019) may be a contributing factor in rising bleaching thresholds. Similarly, intraspecific diversity may also contribute to changes in the bleaching threshold if tolerant genotypes persist in populations and sensitive genotypes are lost. While this would promote reef resistance to bleaching, the ecological consequences of corresponding reductions in genetic diversity remain to be seen.

The effect of intraspecific diversity in bleaching susceptibility on downstream ecological outcomes of individuals with contrasting phenotypes (i.e. bleached vs. not bleached) is not well understood, despite an abundance of data demonstrating that interspecific differences in bleaching susceptibility are predictive of mortality such that species resistant to bleaching have lower mortality than susceptible species (Loya *et al.*, 2001; McClanahan, 2004; Baker *et al.*, 2008; Bahr *et al.*, 2017; Hughes *et al.*, 2017). Kāne□ohe Bay, Hawai□i, located on the northeast coast of O□ahu, is an opportune system for investigating this question of how intraspecific variability in coral responses to thermal bleaching events driven by climate change influence coral survival. The two dominant reef-building coral species in the bay, *Montipora capitata* and *Porites compressa*, both exhibit differences in thermal performance within and between species during bleaching (Grottoli *et al.*, 2006; Cunning *et al.*, 2016; Wall *et al.*, 2019). Differences in symbiont associations and nutritional plasticity are hypothesized to influence bleaching resistance and resilience between these two species (Grottoli *et al.*, 2006; Cunning *et al.*, 2016; Wall *et al.*, 2019), while the species of symbionts associated with *M. capitata* contributes to differences in thermal tolerance (Cunning *et al.*, 2016). However, the influence of intraspecific variation in bleaching susceptibility on ecological outcomes following bleaching events remains unexplored. Understanding how intraspecific variation influences differential outcomes following thermal stress are critical for understanding how the increasing frequency and severity of coral bleaching events will impact the function of these important ecosystems.

In order to better understand how differences in individual colony responses to thermal stress influences ecological outcomes following a coral bleaching event, we conducted a 2-year monitoring study to characterize the recovery dynamics of bleaching susceptible and bleaching resistant coral colonies of *M. capitata* and *P. compressa* following a regional bleaching event in Kāne□ohe Bay in 2015. Hawai□i experienced anomalously high seawater temperatures in late summer (August – September) of 2015, resulting in widespread coral bleaching throughout the region (Bahr *et al.*, 2017). This was the first consecutive coral bleaching event ever observed on Hawaiian reefs, occurring one year following a regional bleaching event in 2014 (Bahr *et al.*, 2015). A total of 22 conspecific pairs of adjacent corals with contrasting bleaching phenotypes (bleaching susceptible or bleaching resistant) of each species were tagged and georeferenced at each of two reefs in the bay with different environmental conditions. Coral pigmentation recovery and mortality were monitored periodically over the following two years to examine how bleaching influences the ecological trajectories within and between species. These colonies are maintained on a regular basis for the purpose of establishing these corals as a living library archived *in situ* for use in future research on coral bleaching mechanisms, thermal physiology, and climate change resilience.

## Materials and Methods

### Site selection and characterization

This study was initiated during the peak of the 2015 coral bleaching event in Kāne□ohe Bay, O□ahu. The lagoon of Kāne□ohe Bay consists of a shallow network of fringing and patch reefs protected by a barrier reef (Bahr *et al.*, 2015), which restricts seawater exchange with the open ocean leading to regions within the bay with high and low seawater residence times (Lowe *et al.*, 2009). Two patch reefs dominated by the reef-building corals *Montipora capitata* and *Porites compressa* from two different regions of the bay were selected for this study: 1) Inner Lagoon region: patch reef 4 (PR4; 21.4339°N, 157.7984°W), an inshore reef with relatively high terrestrial influence (e.g. sedimentation, fresh water and nutrient run-off) and long (>30 days) seawater residence time (Lowe *et al.*, 2009) and 2) Outer Lagoon region: patch reef 13 (PR13; 21.4515°N, 157.7966°W), a seaward reef with less terrestrial influence and short (<24 hour) seawater residence time (Lowe *et al.*, 2009); Figure 1A,B). Temperature loggers (Onset U22 Hobo; 0.21°C accuracy; 0.02°C resolution) were deployed on the benthos of each reef at ~2 m depth from October 2015 through October 2016, and seawater temperature was recorded in 15-minute intervals. The loggers were cross calibrated in an aquarium prior to deployment. Because these loggers were not deployed until after the peak of the heatwave, data from nearby monitoring stations within each region were used to determine if there were any differences in cumulative heat stress between the two regions. These data were collected from PR12, located ~100 m from PR13 in Outer Lagoon (Figure 1D; Table 1; data courtesy of the Division of Aquatic Resources of the State of Hawaii^1^), and PR1 (Coconut Island; data from PACIOOS^2^), in the same Inner Lagoon flow regime as PR4. Degree heating weeks (DHW) were calculated for the inner and outer lagoon reefs from the PR1 and PR12 data, respectively, according to the methods described by (Liu *et al.*, 2013).

**Figure 1.**
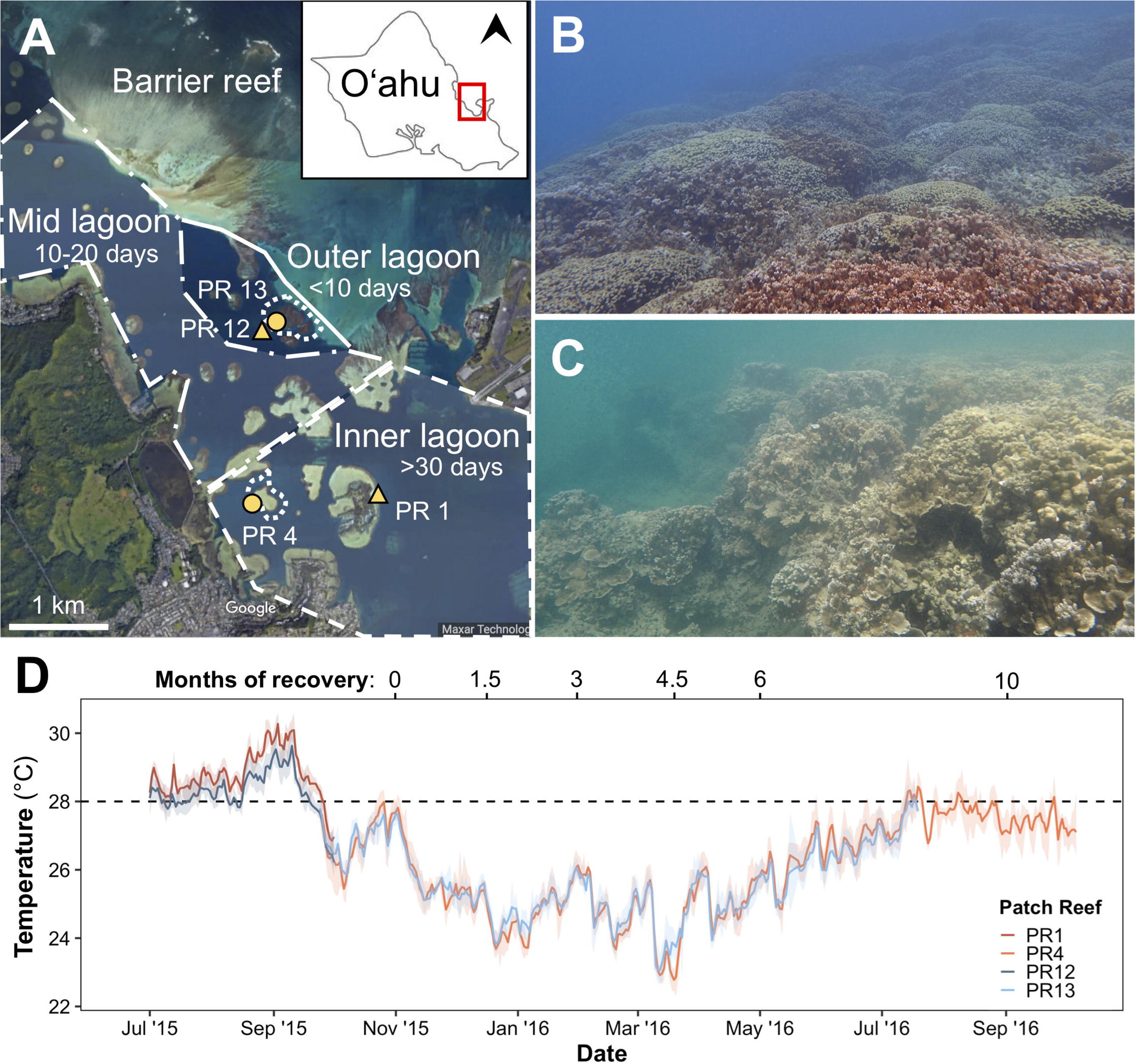
A) Map of the southern region of Kāne□ohe Bay, O□ahu. Inset shows the island of O□ahu, with north indicated by the arrowhead and the red square indicating the southern region of Kāne□ohe Bay enlarged in detail. Distinct hydrodynamic regimes within the lagoon are indicated by the polygons: the dashed line surrounds the Inner Lagoon region where seawater residence times are >30 days; the dash-dot line surrounds the Mid Lagoon region where seawater residence times are 10-20 days; the solid line surrounds the Outer Lagoon region where seawater residence times are <10 days (from Lowe *et al.*, 2009). Yellow symbols indicate locations of *in situ* temperature loggers. Representative images of each reef are shown for B) Outer lagoon (PR13) and C) Inner lagoon (PR4). D) Mean daily temperature at each reef. Shading indicates daily temperature range. Dashed line indicates local coral bleaching threshold.

**Table 1.** Summary of seawater hydrodynamics and temperature conditions within the Inner and Outer lagoon regions in Kāne□ohe Bay, Hawai□i during the peak of the thermal stress event and the initial recovery period. Asterisks indicate statistical significance (p<0.03; Wilcoxon rank sum test).

|  | Region: | Inner Lagoon |  | Outer Lagoon |  | References |
| --- | --- | --- | --- | --- | --- | --- |
|  | Patch Reef: | PR 1<br>(Coconut Island) | PR 4 | PR 12 | PR 13 |  |
| **Physical regime** | Dominant physical force | Tide | Tide | Wave | Wave | Bahr et al. 2017; Lowe et al. 2009 |
|  | Seawater residence time | >30 days | >30 days | 1-5 days | <1 day | Lowe et al. 2009 |
| **Thermal stress**<br>(July 1 - Oct 1, 2015) | Mean maximum daily temperature (°C) | **29.08\*** | ND | **28.64\*** | ND | Figure 1 |
|  | Mean minimum daily temperature (°C) | **28.44\*** | ND | **27.94\*** | ND | Figure 1 |
|  | Degree heating weeks (DHW) | 5.84 | ND | 1.77 | ND | Figure 1 |
| **Initial Recovery**<br>(Nov 1, 2015 - Feb 1, 2016) | Mean maximum daily temperature (°C) | 25.60 | 25.59 | 25.35 | 25.62 | Figure 1 |
|  | Mean minimum daily temperature (°C) | 25.01 | 24.80 | 24.75 | 24.93 | Figure 1 |

### Characterization of benthic community composition and coral bleaching responses

Surveys were performed to assess benthic community composition and coral bleaching responses during the 2015 bleaching event and periodically thereafter for two years. Four surveys were conducted at each site for each time point along the reef slope in parallel to the reef crest at a depth of 2 m (± 1 m). Each survey was carried out along a 40 m transect laid parallel to the reef crest. A photo-quadrat (0.33 m^2^) was placed on the benthos and images were taken every 2 m along the transect using an underwater camera (Canon PowerShot or Olympus TG5). Images were analyzed using Coral Point Counts in excel (CPCe; 100 points per image). Corals were identified to species, and bleaching severity (white, pale, or pigmented) was recorded. Bleaching prevalence was calculated as the proportion of live coral affected by bleaching (white or pale). All other organisms were classified into functional groups (turf algae, crustose coralline algae (CCA), macroalgae, sediment and sand, sponges).

### Monitoring of bleaching resistant and susceptible corals

Individual coral colonies of *M. capitata* and *P. compressa* with contrasting bleaching phenotypes were identified at the peak of the bleaching event from late September through the first week of November 2015. Bleaching resistant corals were defined as those that remained pigmented during the bleaching event, while bleaching susceptible corals were defined as those that appeared completely white. All corals were located along the edge of the reef crest (~1 m depth) and down the reef slope (up to ~3 m depth). A plastic cattle tag with a unique ID was attached to each colony, the colony was photographed, and its GPS location was recorded. All tagged colonies were conspecific pairs of colonies with contrasting bleaching phenotypes that were located adjacent to one another on the reef (e.g. Figure 4). A total of 22 pairs (44 colonies) of each species were tagged at each of two reefs (PR4 and PR13; Supplementary Data 1). This paired design eliminated the potential for differences in the local microenvironment experienced by one phenotype but not the other (e.g. light intensity, flow) to confound interpretation of differences in their ecological outcomes. This paired design is also powerful because it can distinguish between mortality related to thermal stress (both phenotypes) and mortality related to symbiont loss (bleached phenotypes). Bleaching recovery (where applicable) and partial mortality of tagged individuals were monitored every ~6 weeks for the first 6 months following the bleaching event, then once every 6 months up to 24 months following the bleaching event. At each time point, colonies were photographed and visual observations of their bleaching status were recorded. Corals in both the benthic surveys and the tagged individuals were given a color score based on their visual color: 1) white (>80% of colony white with no visible pigmentation); 2) pale (>10% colony affected by pigment loss); or 3) fully pigmented (<10% colony with any pale coloration). Partial mortality was quantified in 20% intervals (0 - 100%) from the photographs. The mean color scores and partial mortality for each species were calculated for each bleaching phenotype and site at each time point. For colonies with a missing observation, mortality scores were estimated using the mortality score of the previous time point. This is a conservative estimate of mortality (i.e. minimum possible mortality at that time point), and in most cases mortality scores did not change surrounding the missing observation. Missing observations of color scores were never interpolated across the time series due to the possibility for rapid loss or recovery of pigmentation.

### Statistical analyses

Statistical analyses were conducted in R v. 3.6.1 (R Core Team, 2017). Seawater temperature means were compared using Wilcoxon rank sum tests. Differences in coral bleaching prevalence and severity from the benthic survey data were compared between species and reefs using 1- or 2-way ANOVAs. For the tagged colonies, the effect of reef lagoon and bleaching phenotype (whether a colony was bleached or pigmented in Nov. 2015) on coral pigmentation and tissue mortality were analyzed at 1.5, 3, 4.5 and 6-months post-bleaching using ANOVAs. In addition, the rates of pigmentation recovery and the rates of tissue loss during the first three and six months following bleaching were each compared between bleaching phenotypes and reefs using 2-way ANOVAs. To control for local environment, a 1-way ANOVA was used to test for the effect of lagoon on the differences in pigmentation score and mortality score between bleaching phenotypes within each pair (*M. capitata*: PR13, n=22 pairs; PR4, n=21 pairs; *P. compressa*: PR13, n=20 pairs; PR4, n=21 pairs). For all approaches (all colonies, paired colonies, differences between pairs, and transect data), Tukey honest significant post hoc tests were used to test for significant pairwise differences in these main effects. To test hypotheses about the differential recovery of bleaching susceptible versus bleaching resistant corals, we used a three way ANOVA model to examine how mean pigmentation and mean tissue loss in each species differed by time, lagoon and bleaching history (B2015) as factorial fixed effects (Table 2&3). We also ran models using pigment or tissue loss difference in adjacent coral pairs as the response variable to account for spatial heterogeneity within sites. Coral pairs with missing measurements were excluded from paired analyses, but this had no effect on patterns of statistical significance (Table 2&3 and Supplementary Table1&2). Models for both 3- and 6-month recovery periods were contrasted in both paired and unpaired analyses (Table 1&2).

**Table 2.** Three-way ANOVA results for pigmentation recovery of bleaching susceptible and resistant colonies during the first six months of recovery for *Montipora capitata* and *Porites compressa* at PR13 and PR4.

|  |  | *Months 0-3* |  |  |  | *Months 0-6* |  |  |  |
| --- | --- | --- | --- | --- | --- | --- | --- | --- | --- |
|  |  | Df | Sum Sq | F val | p | Df | Sum Sq | F val | p |
| *Montipora capitata* | Month | 2 | 14.31 | 68.97 | **<0.01** | 4 | 20.55 | 41.04 | **<0.01** |
|  | Lagoon | 1 | 0.05 | 0.51 | 0.48 | 1 | 0.24 | 1.89 | 0.17 |
|  | B2015 | 1 | 100.16 | 965.73 | **<0.01** | 1 | 101.97 | 814.53 | **<0.01** |
|  | Month:Lagoon | 2 | 0.36 | 1.74 | 0.18 | 4 | 1.21 | 2.42 | **<0.05** |
|  | Month:B2015 | 2 | 11.92 | 57.49 | **<0.01** | 4 | 22.85 | 45.63 | **<0.01** |
|  | Lagoon:B2015 | 1 | 0.05 | 0.49 | 0.49 | 1 | 0.91 | 7.27 | **<0.01** |
|  | Month:Lagoon:B2015 | 2 | 0.76 | 3.67 | **0.03** | 4 | 1.54 | 3.09 | **0.02** |
|  | Residuals | 221 | 22.92 |  |  |  | 351 | 43.94 |  |
| *Porites compressa* | Month | 2 | 46.55 | 196.59 | **<0.01** | 4 | 67.20 | 196.45 | **<0.01** |
|  | Lagoon | 1 | 0.48 | 4.01 | **<0.05** | 1 | 0.68 | 8.00 | **<0.01** |
|  | B2015 | 1 | 34.50 | 291.46 | **<0.01** | 1 | 20.91 | 244.52 | **<0.01** |
|  | Month:Lagoon | 2 | 0.33 | 1.39 | 0.25 | 4 | 0.55 | 1.60 | 0.17 |
|  | Month:B2015 | 2 | 35.49 | 149.89 | **<0.01** | 4 | 49.10 | 143.54 | **<0.01** |
|  | Lagoon:B2015 | 1 | 0.03 | 0.28 | 0.60 | 1 | 0.02 | 0.26 | 0.61 |
|  | Month:Lagoon:B2015 | 2 | 1.63 | 6.87 | **<0.01** | 4 | 1.64 | 4.79 | **<0.01** |
|  | Residuals | 228 | 26.99 |  |  | 375 | 32.07 |  |  |

**Table 3.** Three-way ANOVA results for partial mortality over the first six months post peak bleaching for bleaching susceptible and resistant colonies of *Montipora capitata* and *Porites compressa* at PR13 and PR4.

|  |  | *Months 0-3* |  |  |  | *Months 0-6* |  |  |  |
| --- | --- | --- | --- | --- | --- | --- | --- | --- | --- |
|  |  | Df | Sum Sq | F val | p | Df | Sum Sq | F val | p |
| *Montipora capitata* | Month | 2 | 1.06 | 18.15 | **<0.01** | 4 | 2.63 | 18.26 | **<0.01** |
|  | Lagoon | 1 | 0.72 | 25.15 | **<0.01** | 1 | 2.55 | 70.88 | **<0.01** |
|  | B2015 | 1 | 1.18 | 41.18 | **<0.01** | 1 | 2.93 | 81.49 | **<0.01** |
|  | Month:Lagoon | 2 | 0.43 | 7.39 | **<0.01** | 4 | 0.97 | 6.75 | **<0.01** |
|  | Month:B2015 | 2 | 0.40 | 6.93 | **<0.01** | 4 | 0.63 | 4.39 | **<0.01** |
|  | Lagoon:B2015 | 1 | 0.27 | 9.33 | **<0.01** | 1 | 0.53 | 14.58 | **<0.01** |
|  | Month:Lagoon:B2015 | 2 | 0.15 | 2.59 | 0.08 | 4 | 0.17 | 1.15 | 0.33 |
|  | Residuals | 237 | 6.81 |  |  | 370 | 13.32 |  |  |
| *Porites compressa* | Month | 2 | 0.08 | 2.63 | 0.07 | 4 | 0.22 | 2.87 | **0.02** |
|  | Lagoon | 1 | 0.20 | 13.35 | **<0.01** | 1 | 0.11 | 5.73 | **0.02** |
|  | B2015 | 1 | 0.65 | 44.02 | **<0.01** | 1 | 1.36 | 72.17 | **<0.01** |
|  | Month:Lagoon | 2 | 0.01 | 0.48 | 0.62 | 4 | 0.11 | 1.47 | 0.21 |
|  | Month:B2015 | 2 | 0.05 | 1.63 | 0.20 | 4 | 0.08 | 1.02 | 0.40 |
|  | Lagoon:B2015 | 1 | 0.27 | 18.24 | **<0.01** | 1 | 0.35 | 18.41 | **<0.01** |
|  | Month:Lagoon:B2015 | 2 | 0.03 | 0.96 | 0.39 | 4 | 0.04 | 0.51 | 0.73 |
|  | Residuals | 234 | 3.43 |  |  | 385 | 7.28 |  |  |

## Results

### Temperature dynamics throughout bleaching and recovery

Seawater temperatures exceeded 30°C across Kāne□ohe Bay in late September 2015 (Figure 1D; (Bahr *et al.*, 2017). The average maximum daily seawater temperature during the heatwave was 0.44°C warmer in the Inner Lagoon (29.08°C) than the Outer Lagoon (28.64°C, Table 1). This resulted in higher accumulated heat stress, with a total of 5.84 degree heating weeks (DHW) at Inner Lagoon versus 1.77 DHW at the Outer Lagoon (Table 1). During the recovery period following the heatwave, temperature dynamics were not significantly different between the two regions (Figure 1D; Table 1).

### Benthic community composition

Coral cover was significantly higher at PR13 in the Outer Lagoon (80% ± 4.2%; Figure 2B) than at PR4 in the Inner Lagoon (52% ± 6.0%; Figure 2A) at the initiation of this study (post hoc p=0.01, Supplementary Table 3). Coral cover did not change significantly at either site over the course of the two year bleaching recovery period (Figure 2; Supplementary Table 4). The coral community at PR13 was dominated by *P. compressa* throughout the time series (62-77% of live coral cover) relative to *M. capitata* (23-38% of live coral cover). The relative abundance of each species at PR4 was 54-63% for *P. compressa* vs. 37-46% for *M. capitata*. Turf algae were the second most abundant functional group at both reefs, comprising 6-10% of the benthos at PR13 and 25-40% at PR4. Sediments were also common along the benthos at PR4 (8.5-21%; Figure 2A), but were found in low abundance at PR13 (<1%; Figure 2B). Both reefs had a low abundance (<5%) of macroalgae, crustose coralline algae (CCA) and sponges. PR4 had an abundance of small filter feeders (e.g. boring oysters and sponges) that were not readily apparent on photo-quadrat images but were observed by divers/snorkelers during surveys. This reef also typically had low visibility (<5 m; Figure 1C), whereas PR13 tended to have higher visibility (~20-30 m, personal observations; Figure 1B).

**Figure 2.**
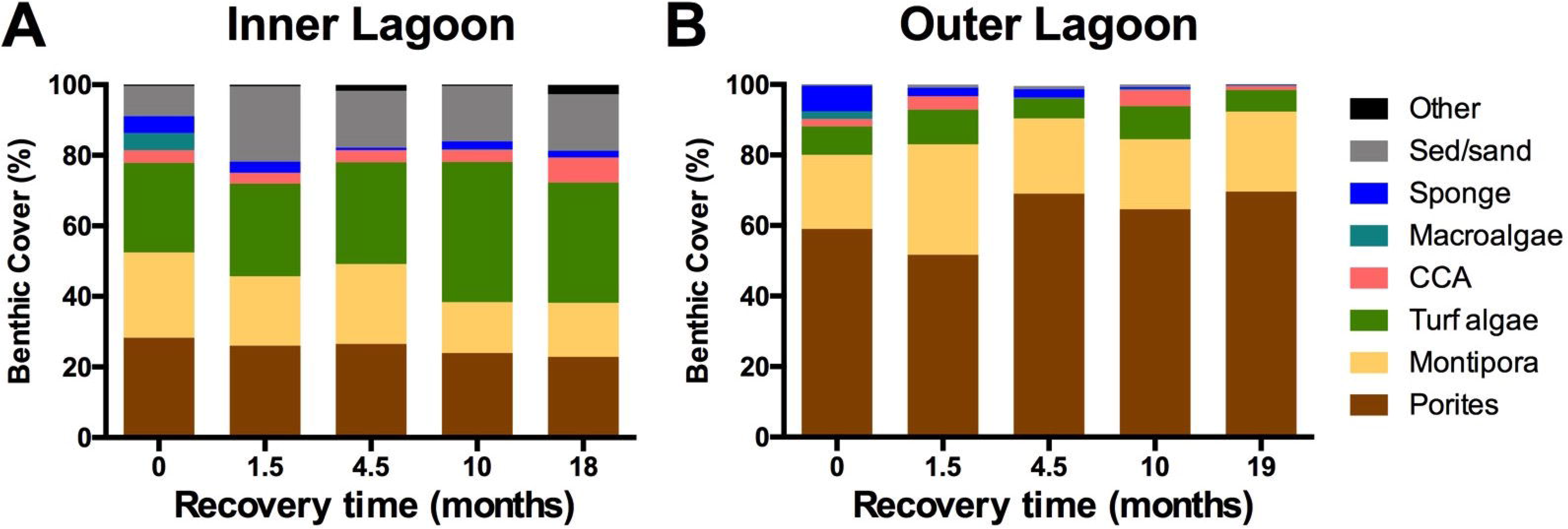
Benthic community composition from the peak of the 2015 bleaching event through 18 months of recovery for A) the Inner lagoon (PR4) and B) the Outer lagoon (PR13) in Kāne□ohe Bay, O□ahu. Data represent the means of four replicate transects.

### Initial coral bleaching prevalence and severity differed between species and reefs

Coral tagging and benthic surveys were initiated during the peak extent of bleaching in Kāne□ohe Bay in late September through October 2015 (Bahr *et al.*, 2017). At that time, there was a higher prevalence of bleaching and paling corals (proportion of live coral that was white or pale) for both species at PR4 in the Inner Lagoon: 69% ± 3% at PR4 vs. 39% ± 12% at PR13 for *M. capitata*; 87% ± 7% at PR4 vs. 45% ± 5% at PR13 for *P. compressa* (post hoc p<0.05; Figure 3; Supplementary Table 5). Within reefs, the prevalence of completely bleached (white) tissue was lower for *M. capitata* (26% ± 6% of all live tissue) than for *P. compressa* (71% ± 10% of all live tissue) at PR4 (p<0.01; Supplementary Table 6; Supplementary Figure 1), whereas there was no significant difference between species at PR13 (Figure 3; Supplementary Table 6). Bleaching severity (the proportion of affected tissue that was completely bleached) was higher for *P. compressa* at PR4 (80% ± 5%) than for *M. capitata* (37% ± 7%, Figure 3; p<0.01, Supplementary Table 7). This high level of bleaching severity for *P. compressa* at PR4 was also higher than this species suffered at PR13 (29% ± 10%; post hoc p<0.01; Supplementary Table 7), whereas bleaching severity did not differ significantly between sites for *M. capitata* (33% ± 5% at PR13 vs. 37% ± 7% at PR4; p>0.05; Supplementary Table 8).

**Figure 3.**
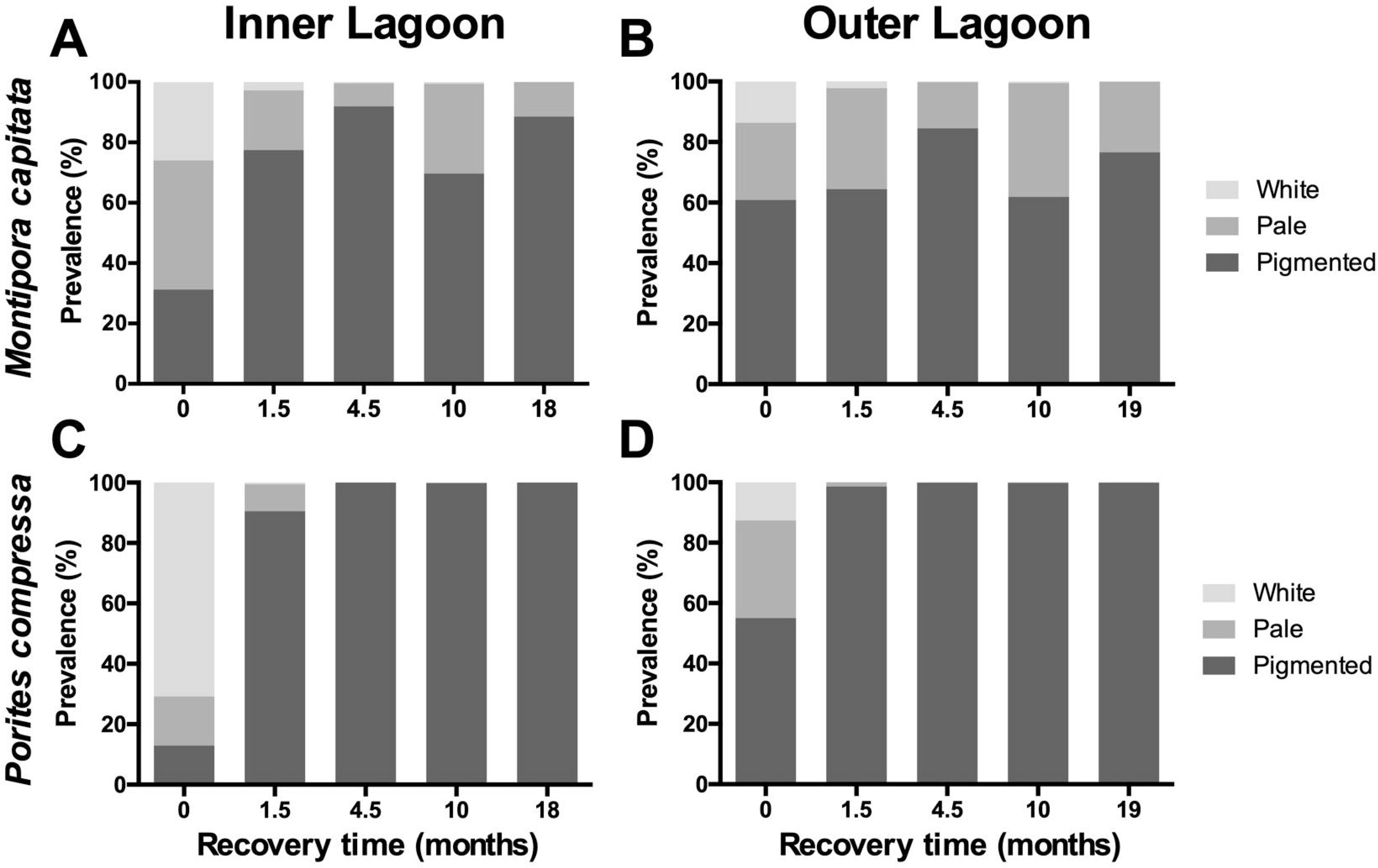
Prevalence of coral bleaching phenotypes for *Montipora capitata* (A-B) and *Porites compressa* (C-D) at the Inner lagoon (left column) and Outer lagoon (right column). Data represent the means of four replicate transects.

### Coral bleaching prevalence differs between species and reefs throughout recovery

Bleaching prevalence rapidly decreased for *P. compressa* at both reefs, with a 97% decrease in bleaching prevalence observed after 1.5 months of recovery at PR13 to <2% overall prevalence, and a 89% decrease at PR4 to <10% prevalence (Figure 3). In contrast, bleaching prevalence in *M. capitata* declined more slowly, with a mean decrease of 67% at the PR4 versus only a 9% decrease at PR13 after 1.5 months. At that point, <3% of *M. capitata* remained fully bleached at either site, and 20-33% remained pale. In contrast to peak bleaching, coral bleaching prevalence was significantly higher for *M. capitata* than *P. compressa* at each time point in the first year of recovery (p<0.01; Supplementary Table 9). Initial differences in bleaching prevalence between reefs at the peak of bleaching also disappeared beginning at 1.5 months of recovery for *M. capitata* and 3 months of recovery for *P. compressa* (p>0.05, Supplementary Table 10; Figure 3). Interestingly, after several months of declining prevalence, the seasonal peak in water temperatures in September 2016 (Month 10; Figure 1D) corresponded with an increase in the prevalence of pale *M. capitata* at both reefs (Figure 3A,B). *P. compressa* pigmentation did not appear affected by this seasonal warming, and bleaching prevalence remained below 2% from 1.5 months of recovery onward (Figure 3C,D).

### Pigmentation recovery dynamics of individual coral colonies following thermal stress

Coral pigmentation increased significantly over time for bleached colonies of both *M. capitata* and *P. compressa*, (p < 0.01; Supplementary Table 11), while bleaching resistant individuals remained pigmented throughout the entire 24 month recovery period (Figures 4&5). The difference in color scores between bleaching susceptible and resistant coral pairs shrank significantly over time (p<0.01; Table 2), indicating significant recovery from bleaching. Over both 3- and 6-month recovery periods, the temporal trajectory of susceptible and resistant corals differed significantly between sites in both coral species (p<0.05; Table 2;); for *M. capitata*, there was a larger difference between pairs at PR13 than PR4 during the 3-6 month timepoints, while for *P. compressa* there was not (indicative of a full recovery). The statistical patterns were robust when analyzing paired differences between susceptible and resistant corals as the response variable (Supplementary Table 12; Supplementary Figure 2). Mean pigmentation did not differ between sites in *M. capitata* in the first 6 months (p>0.05), but *P. compressa* exhibited greater overall pigmentation at PR13 than at PR4 (p<0.01; Table 2). The timing of pigmentation recovery of the bleaching susceptible colonies differed between species. Bleaching susceptible colonies of *P. compressa* remained distinguishable from resistant colonies 1.5 months post peak bleaching (p<0.01), but recovered full visual pigmentation by three months at both reefs (p>0.05; Supplementary Table 11; Figure 5C,D). In contrast, recovery was slower for *M. capitata*, which on average did not recover full pigmentation within the first 6 months (p<0.01, Supplementary Table 11). Only 31% of bleaching susceptible *M. capitata* had fully recovered pigmentation at PR4 after 3 months, while none had fully recovered at PR13 (Figure 5A,B), resulting in a significantly smaller mean difference between susceptible and resistant corals at PR4 relative to PR13 at month 3 (p=0.03), which remained so through month 6 of recovery (p=0.02; Supplementary Table 11). There was a decrease in the mean pigmentation score of bleaching susceptible colonies of *M. capitata* in the late summer of 2016 (month 10), mirroring the increased prevalence of pale *M. capitata* observed in the benthic survey data (Figure 5A,B). This seasonal paling was not observed for *P. compressa* colonies of either bleaching susceptibility phenotype (Figure 5C,D). Full pigmentation recovery for bleaching susceptible *M. capitata* colonies took as long as 24 months at PR4, going from 25% of individuals with full pigmentation at 18 months to 94% at 24 months (Figure 5A). At PR13, most bleaching susceptible *M. capitata* colonies (68%) remained pale for the entire 24-month duration of this study (Figure 5B).

**Figure 4.**
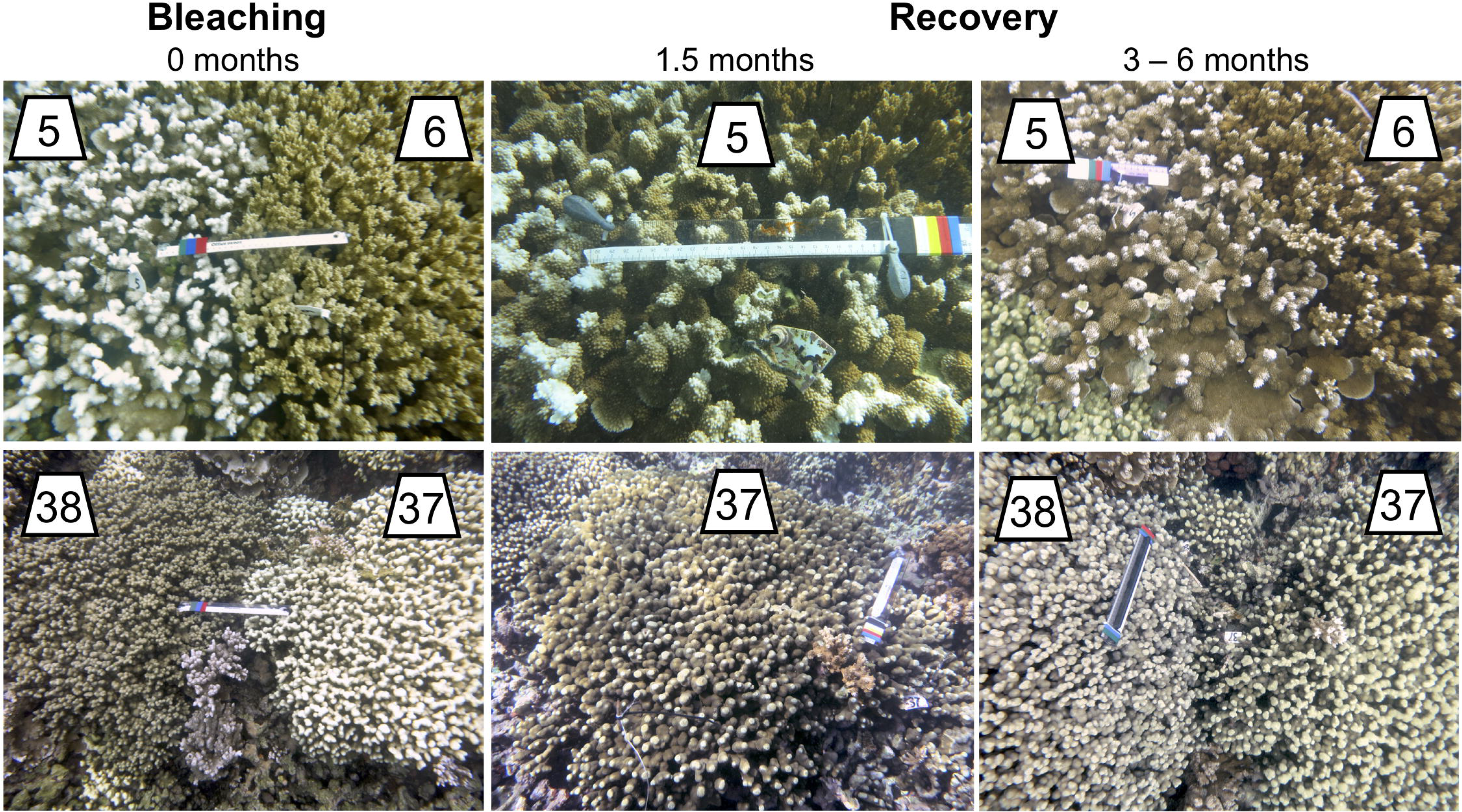
Representative images of tagged bleached and non-bleached corals. Top row: *M. capitata* pair 5&6; Bottom row: *P. compressa* pair 37&38. Left column shows pairs at the peak of the bleaching event in November 2015. Center column shows the bleached colony of the pair after 1.5 months of recovery. Right column shows the same pairs after 3 - 6 months of recovery.

**Figure 5.**
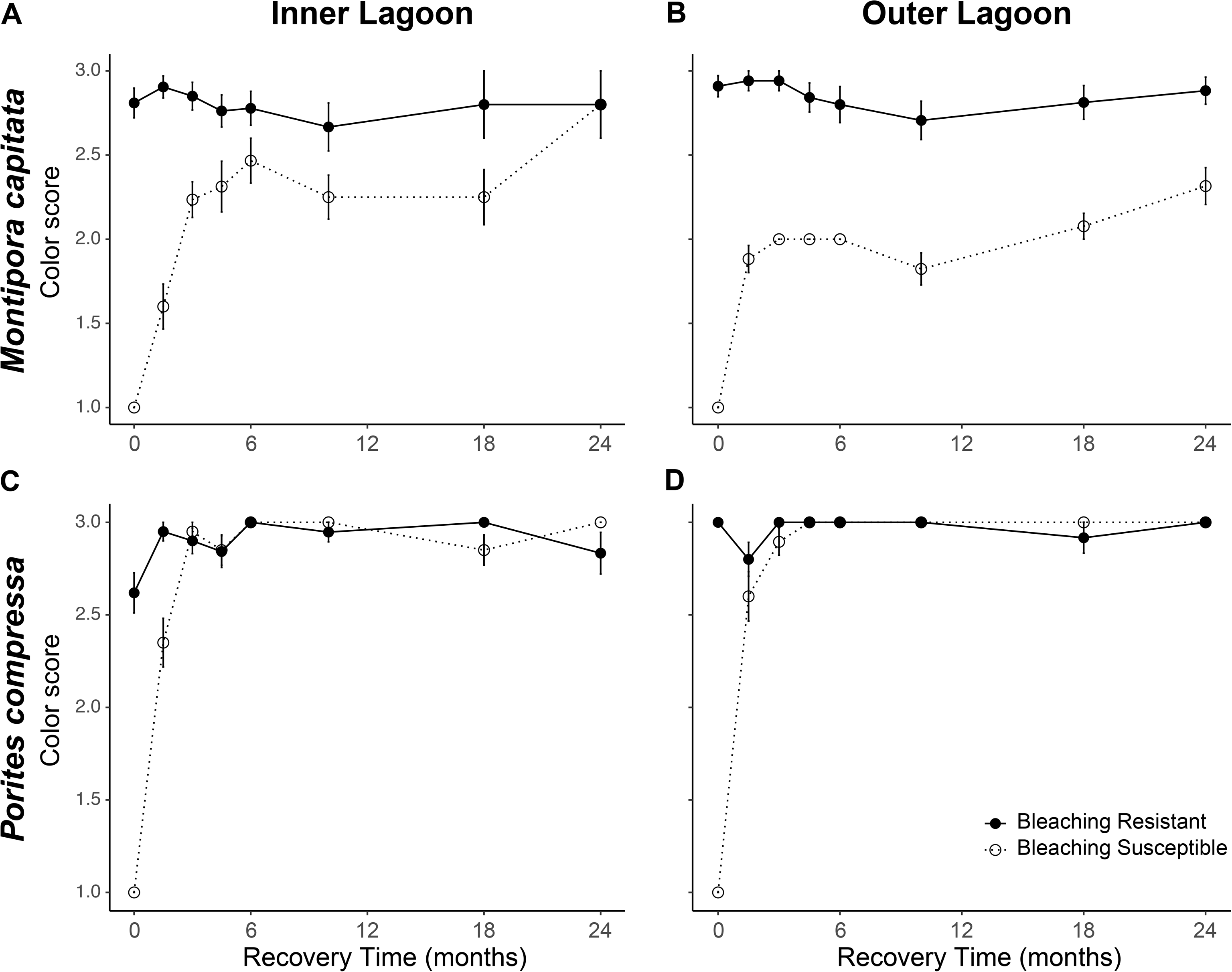
Average color score of bleaching susceptible versus bleaching resistant colonies of *Montipora capitata* (A-B) and *Porites compressa* (C-D) at the Inner Lagoon (A,C) and Outer Lagoon (B,D). Solid lines indicate bleaching resistant colonies; dashed lines indicate bleaching susceptible colonies. Color scores: 1, white; 2, pale; 3, pigmented. Error bars indicate SEM.

### Colony mortality following bleaching differs between bleaching phenotypes

Partial mortality increased significantly for both species in the first 6 months following the bleaching event regardless of bleaching phenotype (p<0.02; Table 3; Figure 6). Bleaching resistant colonies had significantly less cumulative partial mortality than bleaching susceptible colonies across both sites during this same time frame (Table 2; p<0.01). Partial mortality also differed significantly between the two sites, with both species undergoing higher partial mortality at PR4 (p<0.02; Table 2). Differences in tissue loss between susceptible and resistant colonies remained different between species during the first 6 months of recovery (p<0.01; Table 3). The statistical patterns did not differ when analyzing paired differences as the response variable in all cases except one (Supplementary Table 11). Interestingly, in *M. capitata*, during the first 3 months there was no significant effect of the three-way interaction of month, lagoon and bleaching history (p>0.05); however, there was when controlling for location (paired differences) (p=0.03; Supplementary Table 12; Supplementary Figure 2), suggesting spatial location on the reef influences partial mortality following thermal stress.

**Figure 6.**
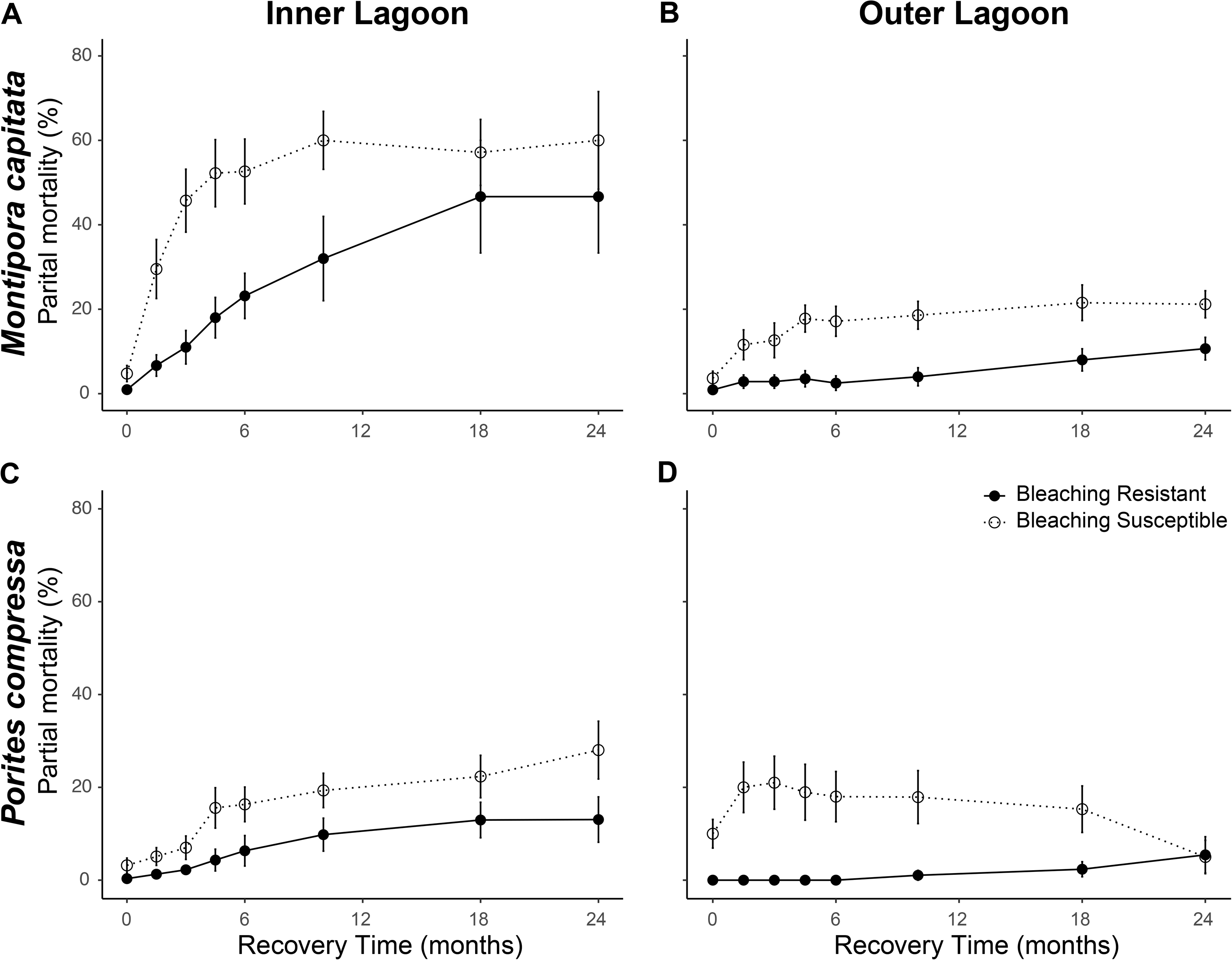
Average partial mortality of bleaching susceptible versus bleaching resistant colonies of *Montipora capitata* (A-B) and *Porites compressa* (C-D) at the Inner Lagoon (A,C) and Outer Lagoon (B,D). Solid lines indicate bleaching resistant colonies; dashed lines indicate bleaching susceptible colonies. Error bars indicate SEM.

Bleaching susceptible colonies of both coral species had higher rates of tissue loss in the first three months following the bleaching event than bleaching resistant corals (post hoc p < 0.01; Supplementary Table 12; Figure 6). Site was also a significant factor for the rate of tissue loss for *M. capitata* but not *P. compressa*, as the rate of tissue loss in the first three months for *M. capitata* was significantly higher at PR4 than PR13 (post hoc p<0.01; Supplementary Table 13; Figure 6A,B). Cumulative partial mortality of bleaching susceptible corals in the first 6 months was relatively low (~17%) for *M. capitata* at PR13 and *P. compressa* at both sites (Figure 6C,D). This was in contrast to the higher partial mortality of both bleaching resistant and bleaching susceptible colonies of *M. capitata* at PR4 (20% and 51% mortality, respectively; Figure 6C,D). Partial mortality in bleaching susceptible colonies of *P. compressa* was significantly greater than bleaching resistant colonies at PR13 in the first three months following bleaching (post hoc p<0.01), after which there was no difference between lagoons (Supplementary Table 12). At three months of recovery, bleaching susceptible *M. capitata* also suffered only ~13% tissue loss at PR13, but had significantly higher partial mortality at PR4 (46% at 3 months (post hoc p < 0.01; Supplementary Table 12). Bleaching susceptible *M. capitata* at PR13 did not differ from bleaching resistant colonies at 3 months (post hoc p = 0.49), whereas there was a difference between phenotypes at PR4 (post hoc p <0.01; Supplementary Table 12). During the two year recovery period, only five colonies (one *P. compressa* and four *M. capitata*) experienced full mortality and all were located at PR4. Tissue loss slowed around 6 months following the bleaching event for bleaching susceptible individuals of both species at both reefs (Figure 6). While bleaching resistant corals showed very low partial mortality in the first 6 months following the heat wave (<25% for both species at both sites; Figure 6), tissue loss gradually increased over the following 24 months, with *M. capitata* at PR4 experiencing the highest loss at 24 months. Of all the corals experiencing partial mortality, *P. compressa* at PR13 was the only group that showed an increase in live tissue within 24 months of the bleaching event (Figure 6D). For example, *P. compressa* (TagID 225; PR13) suffered ~60% mortality in the first two months after undergoing bleaching, but had regrown to nearly 100% live tissue within one year.

## Discussion

### Bleaching susceptibility of a species is not predictive of mortality

*P. compressa* experienced higher bleaching prevalence and severity than *M. capitata* at the inner lagoon reef, and yet bleaching susceptible individuals of *P. compressa* experienced significantly less partial mortality than bleaching susceptible individuals of *M. capitata* at this same site. These results are in contrast to the common pattern of greater bleaching prevalence of a species leading to greater mortality (Baird and Marshall, 2002; Jones, 2008; Hughes, Kerry, *et al.*, 2018), and may reflect two different species-specific thermal stress response strategies. In the case of *M. capitata*, this species resisted bleaching to a greater extent, but individuals that bleached had lower resilience following bleaching (slower recovery and higher mortality). *P. compressa*, on the other hand, was more susceptible to bleaching but had greater resilience following bleaching (faster recovery and lower mortality). The relatively lower resilience of *M. capitata* following bleaching observed here contrasts with experimental predictions that *M. capitata* has a higher capacity to recover from bleaching than *P. compressa* due to its ability to rapidly replace metabolized energy stores by increasing its heterotrophic feeding rates and recover depleted tissue biomass more quickly following bleaching (Grottoli *et al.*, 2006; Rodrigues and Grottoli, 2007). Indeed, bleached *M. capitata* recovered pigmentation and biomass more quickly than *P. compressa* following the 2014 coral bleaching event (Wall *et al.*, 2019), indicating that coral recovery rates following thermal stress may depend on the frequency of thermal stress events. As our observations describe the responses of these corals to the second of two bleaching events to occur within the span of one year, the discrepancies between the outcomes we observed and those of previous studies could be due in part to the recurrent nature of the stress. This pattern has also been observed in other reef systems, where annual repeat bleaching events turn some coral species from winners into losers (Grottoli *et al.*, 2014).

### Ecological outcomes differ between bleaching susceptible and resistant phenotypes

Intraspecific differences in bleaching susceptibility had a significant influence on the ecological outcomes of those individuals in the months following thermal stress. For both *M. capitata* and *P. compressa*, bleaching susceptible individuals suffered higher partial mortality than bleaching resistant individuals located on the same reef. This resulted in significant losses of live coral cover from the reef, which likely has a significant impact on the ecological function of each reef. From an evolutionary perspective, these differences in partial mortality are likely to negatively impact reproductive success, as coral reproductive output is positively correlated with colony size (Hall and Hughes, 1996). Differential partial mortality between susceptible and resistant phenotypes therefore likely influences individual fitness and thus the genetic composition of offspring released in subsequent reproductive events. Further exacerbating this loss, corals that have recently undergone bleaching have a lower likelihood of reproducing at all relative to bleaching resistant conspecifics, and those that do manage to reproduce tend to release fewer and less provisioned gametes (Ward *et al.*, 2000; Fisch *et al.*, 2019). Together these factors will likely limit the evolutionary success of bleaching susceptible genotypes and the ecological resilience of the reef by reducing the recruitment pool and live coral cover (Fisch *et al.*, 2019; Hughes *et al.*, 2019). However, the low frequency of complete colony mortality during this bleaching event indicates the adult gene pool has remained mostly intact, maintaining the genetic diversity of the current population. Intraspecific differences in coral mortality also affect the ecological landscape of coral symbionts. In *M. capitata*, for example, bleaching resistant phenotypes tend to be dominated by *Durusdinium glynii*, whereas bleaching susceptible individuals are mostly dominated by *Cladocopium* sp. (Cunning *et al.*, 2016), and higher survival of *D. glynii*-dominated individuals likely increases the relative abundance of *D. glynii* in the community. As symbiont transmission occurs vertically in this species (Padilla-Gamiño *et al.*, 2012), this would also increase the proportion of larvae inheriting *D. glynii*, potentially altering the composition of the symbiont community for generations. Furthermore, if thermal stress events increase in frequency and severity as predicted (Hughes, Anderson, *et al.*, 2018), the relative growth benefits associated with hosting thermally sensitive symbionts like *Cladocopium* spp. at non-stressful temperatures may fail to make up for the higher costs of these associations during thermal stress, altering the tradeoffs of the association (Cunning *et al.*, 2015). Mechanisms driving differential thermal performance in *P. compressa* are less well understood, particularly as symbiont variation in this species is limited (LaJeunesse *et al.*, 2004).

### Coral bleaching and recovery dynamics differed between inner and outer lagoon reefs

*M. capitata* and *P. compressa* both experienced higher bleaching prevalence and severity at the inner lagoon reef, which corresponded with higher partial mortality for *M. capitata.* These results are reflective of coral mortality patterns from baywide surveys, which also observed higher cumulative coral mortality on reefs in the inner lagoon region (Bahr *et al.*, 2017). Higher rates of bleaching and mortality within the inner lagoon were likely due to the higher accumulated thermal stress at this site. However, additional environmental factors were likely involved because bleaching resistant *M. capitata* also had higher mortality at the inner lagoon reef relative to either phenotype at the outer lagoon reef. These data suggest that differences in coral bleaching and mortality between the two reefs were likely due to a combination of environmental factors, with worse outcomes within the inner lagoon potentially driven by the lower flow rates, longer seawater residence times (>30 days; (Lowe *et al.*, 2009), and closer proximity to land relative to the outer lagoon reef. Low flow environments can limit nutrient and waste exchange in the coral boundary layer, suppressing coral metabolism (Mass *et al.*, 2010; Putnam *et al.*, 2017), and when coupled with higher freshwater and allochthonous nutrient input (Bahr *et al.*, 2015) may have exacerbated the stress of the heat wave, limiting coral recovery and exacerbating tissue loss. Indeed, pigmentation recovery rates were slower within the inner lagoon, a recovery pattern that was also observed in 2014 (Cunning *et al.*, 2016; Wall *et al.*, 2019).

### Coral recovery requires more time between bleaching events

Consecutive annual bleaching events have become a feature of coral reefs around the world (Grottoli *et al.*, 2014; Bahr *et al.*, 2017; Hughes, Anderson, *et al.*, 2018). Short intervals between thermal stress events that prevent individual corals from fully recovering energetically from the first thermal stress prior to exposure to the second are likely to make individuals less resistant to bleaching and mortality in the second event. The 2015 bleaching event followed here occurred one year following the previous bleaching event in 2014, and led to higher cumulative coral mortality than either of the two previous bleaching events in the bay (Bahr *et al.*, 2017), which may have been due in part to the short recovery interval between the two thermal stress events. Indeed, colonies in Kāne□ohe Bay that bleached in 2014 tended to bleach again in 2015 (Ritson-Williams, 2017), suggesting that these corals had not acclimatized to higher temperatures and were unlikely to have fully recovered from that event when the 2015 event occurred. Longer recovery periods are also important for regrowth of tissue lost during a heatwave, and our data showed that tissue lost due to partial mortality was not replaced by live coral cover in the two years following the bleaching event. This indicates that corals in Kāne□ohe Bay require longer recovery intervals to replace lost coral cover following bleaching-related mortality. Furthermore, our data indicate that symbiont recovery rates may take longer following repeat bleaching events, as *M. capitata* recovered visual pigmentation and symbiont abundance in a span of 1-3 months in previous bleaching events (Jokiel and Brown, 2004; Cunning *et al.*, 2016; Ritson-Williams, 2017; Wall *et al.*, 2019), while in 2015 we found that it took as much as two years for many bleached *M. capitata* individuals to fully recover pigmentation. In addition, repetitive bleaching events may influence the differential success of the major reef building species in Kāne□ohe Bay. We found that *P. compressa* recovered pigmentation faster than *M. capitata* and had a lower rate of tissue loss. This was similar to patterns of pigmentation recovery and mortality in these same species at other patch reefs in Kāne□ohe Bay during both the 2014 and 2015 bleaching events (Ritson-Williams, 2017), and suggests that *P. compressa* may gain ecological advantage if bleaching events become more common. Encouragingly, both the 2014 and 2015 events led to low rates of complete colony mortality, with <3% of corals suffering 100% mortality (*M. capitata* and *P. compressa;* this study) versus <2% of *M. capitata* in 2014 (Cunning *et al.*, 2015). A further testament to the resilience of corals in this system, there were several colonies that underwent a complete recovery of live tissue cover following significant (>50%) partial mortality, suggesting that some species are highly resilient and that given a few years of recovery between stress events can rapidly replace lost tissue when mortality is incomplete.

### Individual tracking uncovers low levels of partial mortality following thermal stress

Both bleaching susceptible and bleaching resistant individuals suffered partial mortality in the months following the bleaching event, indicating that although resistant individuals did not visually bleach, all corals were negatively affected by thermal stress. Interestingly, while bleaching-susceptible colonies of both species suffered an average of ~20% tissue loss in the first six months following the bleaching event, these losses did not manifest as significant changes in live coral cover. This discrepancy could be due in part to the low mortality of bleaching resistant phenotypes masking the higher mortality of bleaching-susceptible individuals in community wide surveys. In addition, the high variance of photoquadrat surveys makes it difficult to detect small changes in benthic cover (Jokiel *et al.*, 2015). Photoquadrat surveys are also commonly used to quantify recently dead coral cover during or following a bleaching event, however this method cannot discern whether the coral that died had in fact bleached, whereas individual colony data revealed the consequences of thermal stress for both bleaching phenotypes. While these losses in live coral cover were not detected at the community level, the significant partial mortality observed at the colony level has a negative impact on the ecology and evolution of coral communities as described above. Loss of susceptible individuals may also lower the genotypic richness of the population, which in corals can correlate with lower reproductive success (Baums *et al.*, 2013). That said, the lack of a significant decline in coral cover at these two reefs is encouraging for reef recovery from this bleaching event.

### Living library: coral pairs are a resource for future research and restoration

By identifying individual coral colonies with distinct thermal stress responses and tracking them over a multi-year period, we have generated and continue to maintain a live geo-referenced biological archive in the field as a resource for research on the effects of environmental history and thermal stress responses on the ecology and physiology of reef-building corals. This resource will be particularly valuable in light of predicted increases in the frequency and severity of coral bleaching events, as it allows for prospective sampling of bleaching susceptible vs. resistant phenotypes before, during, and following a bleaching event. This experimental system allows researchers to address questions of coral acclimatization and adaptation to changing oceans, such as examining whether individual coral responses predict tolerance to future stress. In addition, it allows researchers to identify if repeat events change the ecological outcomes observed in the current event, potentially altering which species are considered the ecological winners. Finally, fully recovered corals of known contrasting thermal tolerances can be used to investigate the interaction of temperature tolerance with coral responses to other stressors, including but not limited to ocean acidification, eutrophication, and pathogens, helping to identify potential tradeoffs of thermal tolerance in corals and the importance of phenotypic and genetic diversity in coral reef resilience (Stachowicz *et al.*, 2007; Ladd *et al.*, 2017). This will be useful for informing coral restoration strategies, particularly in light of the movement towards human assisted evolution (e.g. assisted gene flow), as it allows assessment of the potential tradeoffs of propagating thermally tolerant genotypes (Van Oppen *et al.*, 2015, 2017). From a management perspective, bleaching resistant individuals have the potential to serve as a resource for reef managers interested in propagating climate change resilient genotypes on damaged or degraded coral reefs (Van Oppen *et al.*, 2017). Comparing bleaching responses within species demonstrates the importance of understanding individual colony susceptibility to thermal stress and trajectories of recovery in the face of ongoing climate change. These questions are important for understanding the persistence of coral reefs into the future.

## Supporting information

Supplemental Datasheet1

## Acknowledgements

We thank Yanitza Grantcharska and George Davies for help in the field, and the Hawaii Institute of Marine Biology staff for logistical support. Thank you also to the State of Hawaii Division of Aquatic Resources for sharing data from their temperature sensors. This is HIMB contribution # and SOEST contribution #.

## Author contributions

The study was designed by KB and RG. EL, JD and JH helped collect the data. KB, SM, AP, TI, and AH collected and analyzed the data. The paper was written by KB and SM, and all authors edited and approved the final version of the manuscript.

## Data accessibility

The datasets analyzed for this study and R scripts can be found in GitHub: https://github.com/BurgessShayle/2015-Coral-bleaching-recovery. The raw data supporting the conclusions of this manuscript will be made available by the authors, without undue reservation, to any qualified researcher.

## Footnotes

1 Division of Aquatic Resources, State of Hawaii; http://dlnr.hawaii.gov/dar/

2 http://www.pacioos.hawaii.edu/weather/obs-mokuoloe/

