## Supplemental Tables for "Coral bleaching susceptibility is predictive of subsequent mortality within but not between coral species"

Supplementary Material

### Supplementary Figures

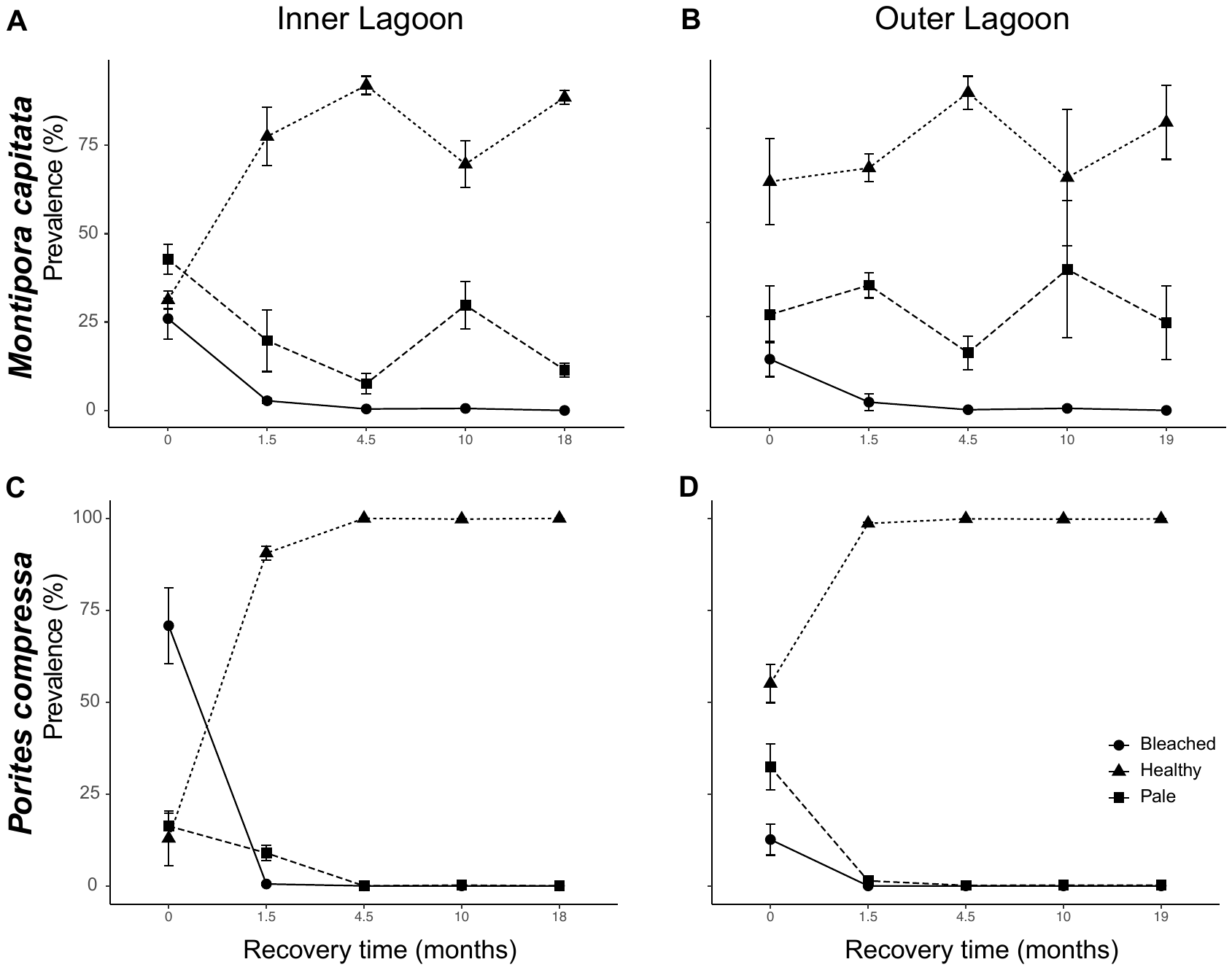

**Supplementary Figure 1.** Prevalence of coral bleaching phenotypes (bleached [circles], healthy [triangles], pale [squares] for *Montipora capitata* (A-B) and *Porites compressa* (C-D) at the Inner lagoon (left column) and Outer lagoon (right column). Data represent the means of four replicate transects. Error bars represent standard error of mean.

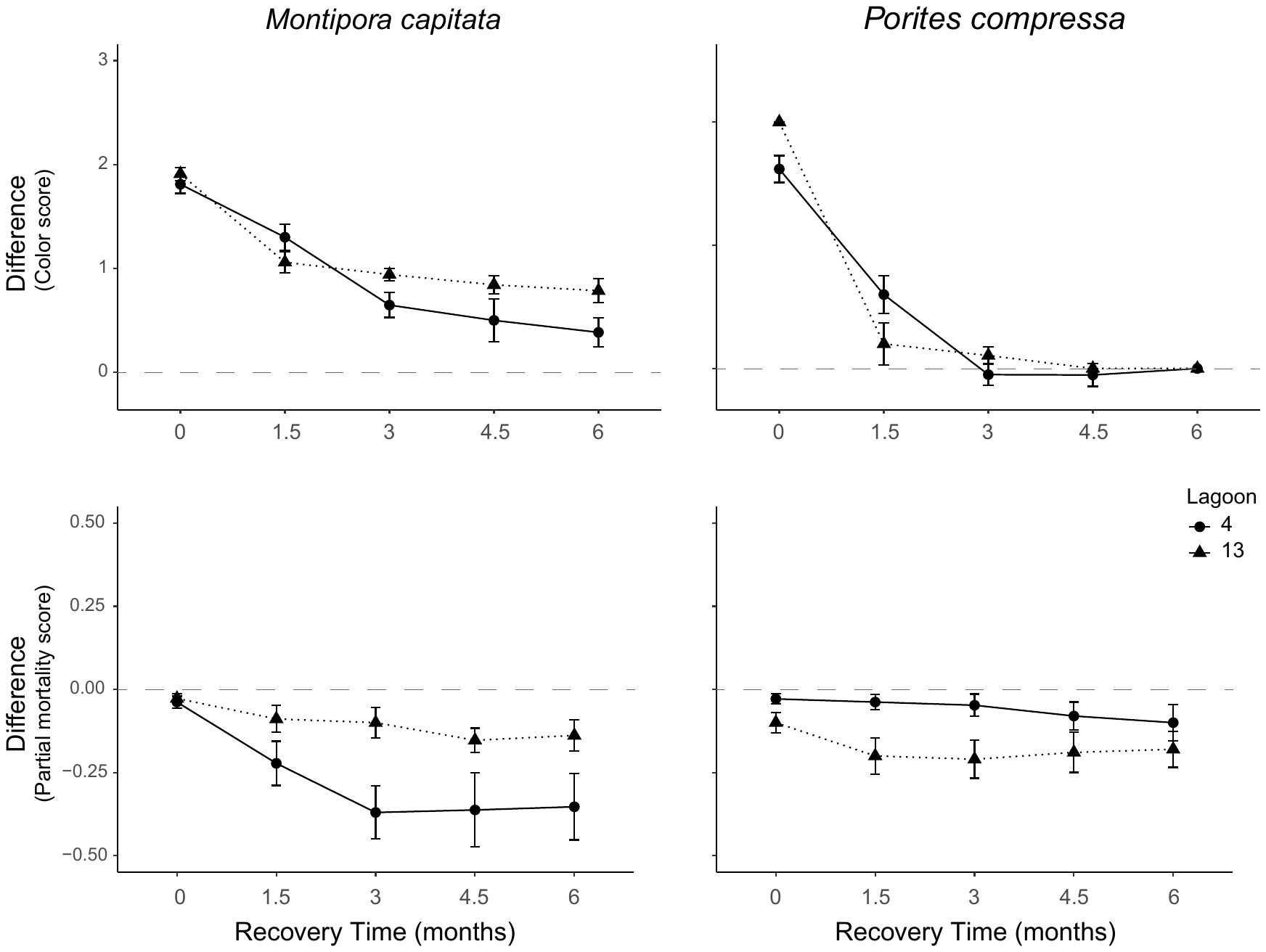

**Supplementary Figure 2.** Mean color (top row) and partial mortality (bottom row) difference scores of adjacent paired colonies (bleaching resistant – bleaching susceptible) of *Montipora capitata* (left column) and *Porites compressa* (right column) at the Inner (PR4, circles) and Outer (PR13, triangles) lagoons. Error bars represent standard error of mean.

### Supplementary Tables

**Supplementary Table 1.** 2-way ANOVA summary table of effects of lagoon and bleaching phenotype (B2015) on pigmentation score of *Montipora* *capitata* and *Porites compressa* during the first 6 months of recovery. Analysis conducted on three datasets: Full – data collected at all time points; Pairs – subset of the Full dataset containing only colonies where both partners of each pair were found; diff – difference between pairs (resistant pigmentation score – susceptible pigmentation score) for each pair. Values indicate p-values. Bold indicates statistical significance.

|  |  | ***Montipora capitata*** | | | ***Porites compressa*** | | |
| --- | --- | --- | --- | --- | --- | --- | --- |
|  |  | Full | Pairs | diff | Full | Pairs | diff |
| Month 0 | Lagoon | 0.36 | 0.34 |  | **<0.01** | **<0.01** |  |
|  | B2015 | **<0.01** | **<0.01** |  | **<0.01** | **<0.01** |  |
|  | Lagoon:B2015 | 0.36 | 0.36 | 0.36 | **<0.01** | **<0.01** | **<0.01** |
| Month 1.5 | Lagoon | 0.09 | 0.13 |  | 0.64 | 0.64 |  |
|  | B2015 | **<0.01** | **<0.01** |  | **<0.01** | **<0.01** |  |
|  | Lagoon:B2015 | 0.2 | 0.19 | 0.16 | 0.07 | 0.07 | 0.09 |
| Month 3 | Lagoon | 0.14 | 0.22 |  | 0.69 | 0.69 |  |
|  | B2015 | **<0.01** | **<0.01** |  | 0.65 | 0.65 |  |
|  | Lagoon:B2015 | **0.03** | **0.05** | **0.03** | 0.17 | 0.17 | 0.18 |
| Month 4.5 | Lagoon | 0.09 | 0.14 |  | **0.02** | **0.03** |  |
|  | B2015 | **<0.01** | **<0.01** |  | 0.95 | 0.64 |  |
|  | Lagoon:B2015 | **0.04** | 0.08 | 0.11 | 0.95 | 0.65 | 0.59 |
| Month 6 | Lagoon | **0.03** | **0.02** |  |  |  |  |
|  | B2015 | **<0.01** | **<0.01** |  |  |  |  |
|  | Lagoon:B2015 | **0.02** | 0.06 | **0.03** |  |  |  |

**Supplementary Table 2.** 2-way ANOVA summary table of effects of lagoon and bleaching phenotype (B2015) on tissue loss score of *Montipora* *capitata* and *Porites compressa* during the first 6 months of recovery. Analysis conducted on three datasets: Full – data collected at all time points; Pairs – subset of the Full dataset containing only colonies where both partners of each pair were found; diff – difference between pairs (resistant pigmentation score – susceptible pigmentation score) for each pair. Values indicate p-values. Bold indicates statistical significance.

|  |  | ***Montipora capitata*** | | | ***Porites compressa*** | | |
| --- | --- | --- | --- | --- | --- | --- | --- |
|  |  | Full | Pairs | diff | Full | Pairs | diff |
| Month 0 | Lagoon | 0.68 | 0.68 |  | **0.04** | **0.04** |  |
|  | B2015 | **0.03** | **0.03** |  | **<0.01** | **<0.01** |  |
|  | Lagoon:B2015 | 0.71 | 0.71 | 0.18 | **0.04** | **0.04** | 0.18 |
| Month 1.5 | Lagoon | **0.01** | **0.01** |  | **0.01** | **0.01** |  |
|  | B2015 | **<0.01** | **0** |  | **<0.01** | **<0.01** |  |
|  | Lagoon:B2015 | 0.1 | 0.11 | 0.63 | **0.01** | **0.01** | 0.63 |
| Month 3 | Lagoon | **<0.01** | **<0.01** |  | 0.06 | 0.06 |  |
|  | B2015 | **<0.01** | **<0.01** |  | **<0.01** | **<0.01** |  |
|  | Lagoon:B2015 | **0.01** | **0.02** | 0.46 | **0.01** | 0.01 | 0.46 |
| Month 4.5 | Lagoon | **<0.01** | **<0.01** |  | **0.89** | **0.72** |  |
|  | B2015 | **<0.01** | **<0.01** |  | **<0.01** | **<0.01** |  |
|  | Lagoon:B2015 | 0.05 | **0.03** | 0.63 | 0.32 | 0.12 | 0.63 |
| Month6 | Lagoon | **<0.01** | **<0.01** |  | 0.59 | 0.59 |  |
|  | B2015 | **<0.01** | **<0.01** |  | **0** | **<0.01** |  |
|  | Lagoon:B2015 | 0.19 | 0.06 | 0.46 | 0.28 | 0.28 | 0.46 |

**Supplementary Table 3.** 1-way ANOVA summary table of the effect of lagoon on coral cover at peak bleaching (Nov 2015). Bold indicates statistical significance. Data collected by benthic transects.

|  | Df | Sum Sq | F value | p |
| --- | --- | --- | --- | --- |
| Lagoon | 1 | 1519.10 | 14.26 | **0.01** |
| Residuals | 6 | 106.30 |  |  |

**Supplementary Table 4.** 1-way ANOVA summary table of the effect of recovery time (months) on coral cover. Data collected by benthic transects.

|  |  | Df | Sum Sq | F value | p |
| --- | --- | --- | --- | --- | --- |
| PR13 | Month | 4 | 426.70 | 1.45 | 0.27 |
|  | Residuals | 15 | 1100.70 |  |  |
| PR4 | Month | 4 | 654.10 | 1.50 | 0.25 |
|  | Residuals | 15 | 1631.30 |  |  |

**Supplementary Table 5.** 1-way ANOVA summary table of the effect of lagoon on bleaching prevalence at peak bleaching (Nov 2015). Bold indicates statistical significance. Data collected by benthic transects.

|  |  | Df | Sum Sq | F value | p |
| --- | --- | --- | --- | --- | --- |
| *Montipora capitata* | Lagoon | 1 | 1751.00 | 6.30 | **<0.05** |
|  | Residuals | 6 | 1668.00 |  |  |
| *Porites compressa* | Lagoon | 1 | 3551.00 | 21.80 | **<0.01** |
|  | Residuals | 6 | 977.00 |  |  |

**Supplementary Table 6.** 1-way ANOVA summary table of the prevalence of bleached (white, not pale) corals at peak bleaching (Nov 2015) by coral species (*Montipora capitata* and *Porites compressa*) at peak bleaching (Nov 2015) at the Inner (PR4) and Outer (PR13) Lagoons. Bold indicates statistical significance. Data collected by benthic transects.

|  |  | Df | Sum Sq | F value | p |
| --- | --- | --- | --- | --- | --- |
| PR4 | Species | 1 | 4022 | 14.46 | **<0.01** |
|  | Residuals | 6 | 1669 |  |  |
| PR13 | Species | 1 | 2.1 | 0.026 | 0.88 |
|  | Residuals | 6 | 478.8 |  |  |

**Supplementary Table 7.** 1-way ANOVA summary table for the effect of coral species on bleaching severity (proportion of completely white corals out of the all affected corals) at peak bleaching (Nov 2015) at PR4. Bold indicates statistical significance. Data collected by benthic transects.

|  | Df | Sum Sq | F value | p |
| --- | --- | --- | --- | --- |
| Species | 1 | 0.37 | 21.71 | **<0.01** |
| Residuals | 6 | 0.1 |  |  |

**Supplementary Table 8.** 1-way ANOVA summary table of the effect of lagoon on bleaching severity of *M. capitata* and *P. compressa* at peak bleaching (Nov 2015). Bold indicates statistical significance. Data collected by benthic transects.

|  |  | Df | Sum Sq | F value | p |
| --- | --- | --- | --- | --- | --- |
| *Montipora capitata* | Lagoon | 1 | 0.00 | 0.23 | 0.65 |
|  | Residuals | 6 | 0.09 |  |  |
| *Porites compressa* | Reef | 1 | 0.53 | 19.37 | **<0.01** |
|  | Residuals | 6 | .16 |  |  |

**Supplementary Table 9.** 1-way ANOVA summary table of the effect of coral species (*Montipora capitata* and *Porites compressa*) on bleaching prevalence (bleached and pale phenotypes). Bold indicates statistical significance. Data collected by benthic transects.

|  |  | Df | Sum Sq | F value | p |
| --- | --- | --- | --- | --- | --- |
| Month 1.5 | Species | 1 | 2223 | 20.93 | **<0.01** |
|  | Residuals | 14 | 1487 |  |  |
| Month 4.5 | Species | 1 | 547.20 | 18.02 | **<0.01** |
|  | Residuals | 14 | 425.10 |  |  |
| Month 10 | Species | 1 | 4624.00 | 14.01 | **<0.01** |
|  | Residuals | 14 | 4622.00 |  |  |

**Supplementary Table 10.** 1-way ANOVA summary table of the effect of reef lagoon (Inner lagoon (PR4) and Outer lagoon (PR13)) on bleaching prevalence (bleached and pale phenotypes) at the Inner (PR4) and Outer (PR13) Lagoons. Bold indicates statistical significance. Data collected by benthic transects.

|  |  | ***Montipora capitata*** | | | | ***Porites compressa*** | | | |
| --- | --- | --- | --- | --- | --- | --- | --- | --- | --- |
|  |  | Df | Sum Sq | F value | p | Df | Sum Sq | F value | p |
| Month 0 | Lagoon | 1 | 1750.60 | 6.30 | **<0.05** | 1 | 3551 | 21.8 | **<0.01** |
|  | Residuals | 6 | 1668.00 |  |  | 6 | 977 |  |  |
| Month 1.5 | Lagoon | 1 | 338.90 | 2.08 | 0.20 | 1 | 130.4 | 18.96 | **<0.01** |
|  | Residuals | 6 | 976.40 |  |  | 6 | 41.27 |  |  |
| Month 4.5 | Lagoon | 1 | 111.90 | 2.14 | 0.19 | 1 | 0.03 | 1.42 | 0.28 |
|  | Residuals | 6 | 313.10 |  |  | 6 | 0.12 |  |  |
| Month 10 | Lagoon | 1 | 121.00 | 0.16 | 0.70 | 1 | <0.01 | **<0.01** | 0.97 |
|  | Residuals | 6 | 4500.00 |  |  | 6 | 0.99 | 0.16 |  |

**Supplementary Table 11.** Two-way ANOVAs testing the effect of lagoon (Inner (PR4) and Outer (PR13) lagoon) and bleaching phenotype (B2015) on *Montipora capitata* and *Porites compressa* color scores over the first 6 months of recovery. Bold indicates statistical significance.

|  |  | ***Montipora capitata*** | | | | ***Porites compressa*** | | | |
| --- | --- | --- | --- | --- | --- | --- | --- | --- | --- |
|  |  | Df | Sum Sq | F value | p | Df | Sum Sq | F value | p |
| Month 0 | Lagoon | 1 | 0.05 | 0.86 | 0.36 | 1 | 0.74 | 11.71 | **<0.01** |
|  | B2015 | 1 | 74.42 | 1206.88 | **<0.01** | 1 | 66.78 | 1051.79 | **<0.01** |
|  | Lagoon:B2015 | 1 | 0.05 | 0.86 | 0.36 | 1 | 0.74 | 11.71 | **<0.01** |
|  | Residuals | 82 | 5.06 |  |  | 78 | 4.95 |  |  |
| Month 1.5 | Lagoon | 1 | 0.48 | 2.98 | 0.09 | 1 | 0.05 | 0.22 | 0.64 |
|  | B2015 | 1 | 26.16 | 1561.98 | **<0.01** | 1 | 3.2 | 13.9 | **<0.01** |
|  | Lagoon:B2015 | 1 | 0.27 | 1.66 | 0.2 | 1 | 0.8 | 3.47 | 0.07 |
|  | Residuals | 70 | 11.31 |  |  | 76 |  |  |  |
| Month 3 | Lagoon | 1 | 0.22 | 2.28 | 0.14 | 1 | 0.01 | 0.16 | 0.69 |
|  | B2015 | 1 | 10.74 | 111.49 | **<0.01** | 1 | 0.01 | 0.21 | 0.65 |
|  | Lagoon:B2015 | 1 | 0.48 | 4.96 | **0.03** | 1 | 0.12 | 1.91 | 0.17 |
|  | Residuals | 68 | 6.55 |  |  | 74 | 4.54 |  |  |
| Month 4.5 | Lagoon | 1 | 0.47 | 2.87 | 0.09 | 1 | 0.44 | 6.2 | **0.02** |
|  | B2015 | 1 | 8.02 | 49.04 | **<0.01** | 1 | 0 | 0 | 0.95 |
|  | Lagoon:B2015 | 1 | 0.73 | 4.43 | **0.04** | 1 | 0 | 0 | 0.95 |
|  | Residuals | 72 | 11.77 |  |  | 71 | 5.08 |  |  |
| Month 6 | Lagoon | 1 | 0.77 | 4.8 | **0.03** | 1 | 0 | 1 | 0.32 |
|  | B2015 | 1 | 4.51 | 28.29 | **<0.01** | 1 | 0 | 1 | 0.32 |
|  | Lagoon:B2015 | 1 | 0.92 | 5.76 | **0.02** | 1 | 0 | 1 | 0.32 |
|  | Residuals | 58 | 9.24 |  |  | 76 | 0 |  |  |

**Supplementary Table 12.** Two-way ANOVAs testing the effect of lagoon (Inner (PR4) and Outer (PR13)) and bleaching phenotype (B2015) on *Montipora capitata* and *Porites compressa* partial mortality over the first 6 months of recovery. Bold indicates statistical significance.

|  |  | ***Montipora capitata*** | | | | ***Porites compressa*** | | | |
| --- | --- | --- | --- | --- | --- | --- | --- | --- | --- |
|  |  | df | Sum Sq | F value | p | df | Sum Sq | F value | p |
| Month 0 | Lagoon | 1 | 0 | 0.17 | 0.68 | 1 | 0.03 | 4.4 | **0.04** |
|  | B2015 | 1 | 0.02 | 5.2 | **0.03** | 1 | 0.08 | 13.89 | **<0.01** |
|  | Lagoon:B2015 | 1 | 0 | 0.14 | 0.71 | 1 | 0.03 | 4.04 | **0.04** |
|  | Residuals | 82 | 0.36 |  |  | 78 | 0.46 |  |  |
| Month 1.5 | Lagoon | 1 | 0.25 | 6.92 | **0.01** | 1 | 0.1 | 6.22 | **0.01** |
|  | B2015 | 1 | 0.52 | 15.32 | **<0.01** | 1 | 0.28 | 16.72 | **<0.01** |
|  | Lagoon:B2015 | 1 | 0.1 | 2.8 | 0.1 | 1 | 0.13 | 7.99 | **0.01** |
|  | Residuals | 78 | 2.84 |  |  | 78 | 1.31 |  |  |
| Month 3 | Lagoon | 1 | 0.92 | 19.56 | **<0.01** | 1 | 0.08 | 3.72 | 0.06 |
|  | B2015 | 1 | 1.02 | 21.64 | **<0.01** | 1 | 0.33 | 15.52 | **<0.01** |
|  | Lagoon:B2015 | 1 | 0.31 | 6.71 | **0.01** | 1 | 0.13 | 6.36 | **0.01** |
|  | Residuals | 77 | 3.61 |  |  | 78 | 1.66 |  |  |
| Month 4.5 | Lagoon | 1 | 0.99 | 21.34 | **<0.01** | 1 | 0 | 0.02 | **0.89** |
|  | B2015 | 1 | 1.11 | 23.74 | **<0.01** | 1 | 0.45 | 15.26 | **<0.01** |
|  | Lagoon:B2015 | 1 | 0.18 | 3.9 | 0.05 | 1 | 0.03 | 1.01 | 0.32 |
|  | Residuals | 69 | 3.21 |  |  | 76 | 2.24 |  |  |
| Month 6 | Lagoon | 1 | 1.37 | 26.55 | **<0.01** | 1 | 0.01 | 0.3 | 0.59 |
|  | B2015 | 1 | 0.89 | 17.35 | **<0.01** | 1 | 0.39 | 14.55 | **<0.01** |
|  | Lagoon:B2015 | 1 | 0.09 | 1.79 | 0.19 | 1 | 0.032 | 1.188 | 0.28 |
|  | Residuals | 64 | 3.30 |  |  | 76 | 2.05 |  |  |

**Supplementary Table 13.** 2-way ANOVA testing the effect of lagoon (Inner (PR4) and Outer (PR13)) and bleaching phenotype (B2015) on the rate of tissue loss during the first three months of recovery. Bold indicates statistical significance.

|  | ***Montipora capitata*** | | | | ***Porites compressa*** | | | |
| --- | --- | --- | --- | --- | --- | --- | --- | --- |
|  | Df | Sum Sq | F value | p | Df | Sum Sq | F value | p |
| Lagoon | 1 | .84 | 21.11 | **<0.01** | 1 | .01 | 1.07 | 0.30 |
| B2015 | 1 | .76 | 19.18 | **<0.01** | 1 | .08 | 6.18 | **0.02** |
| Lagoon:B2015 | 1 | .28 | 6.95 | **0.01** | 1 | .04 | 3.18 | 0.08 |
| Residuals | 77 | 3.06 |  |  | 78 | 1.04 |  |  |
